# Metal speciation in blood plasma

**DOI:** 10.1101/2025.10.04.680475

**Authors:** Apar Prasad, Everett L. Shock

## Abstract

Metal speciation in blood plasma is heavily influenced by proteins and peptides including transferrin, albumin, and glutathione. Despite this, few studies have incorporated these large molecules in speciation calculations, probably due to a lack of experimental measurements. Additionally, there is increasing evidence that metal complexes of small molecules are bioavailable. Due to the limitations posed by analytical techniques, thermodynamic models can serve as an excellent alternative to experimental measurements of metal speciation. In this work, we predict metal speciation for several biologically relevant metals incorporating complexes with proteins, peptides and small molecules. We supplemented experimental measurements from the literature with linear free energy estimates to fulfill the inventory of stability constants required to perform these calculations. In addition to evaluating the speciation of naturally present metals, we also predict the speciation of metals used for therapeutic applications like anticancer drugs, antidiabetics and antacids. Our results indicate that metal speciation is heavily dependent on pH and chelator concentration and can change drastically as metals move from blood plasma to inside cells. Additionally, metal speciation can be dominated by proteins like transferrin and is subject to change as metals cross the blood-brain barrier. Our results corroborate many experimental measurements and can help design future experiments investigating the biological impact of metal-based drugs and metal-toxicity.

## 1. Introduction

Metals are indispensable to life. They serve as cofactors in numerous enzymatic reactions, help stabilize protein conformations, and may also be used as sources of energy via redox transformations. Different metals are present at multiple levels of abundance-while potassium is needed in millimolar concentration in an organism, nickel may be present only at nanomolar levels. Evidently, metal requirements are essential for human life too, iron in hemoglobin of red blood cells being a common example. Over the past 40 years, a mountain of evidence has emerged demonstrating that not all forms of metal are bioavailable (Sunda & Guillard 1976, Anderson & Morel 1978, Sunda & Gillespie 1979, Daly et al. 1990, Canterford & Canterford 1992, Morton et al. 2000, Zhu et al. 2006, Scheers et al. 2014, Peng et al. 2014, Pereira et al. 2014, Nday et al. 2012 and Hart et al. 2015). The organisms investigated in these studies comprise prokaryotes and eukaryotes including human beings. Therefore, quantifying metal distribution (or metal speciation) in a living system can be of enormous benefit, as evidenced by the application of iron speciation in the treatment of iron-overloading (Temraz et al. 2014 and Templeton 2015). However, obtaining metal speciation in complex aqueous systems like human bodily fluids presents extreme experimental difficulties (Levina et al. 2017). Theoretical modeling offers a promising alternative but is generally limited by lack of thermodynamic data (Wilke et al. 2017 and Kiss et al. 2017). Existing models of metal speciation in blood plasma suffer from exclusion of large molecules like albumin and rarely include comparisons with experimental measurements, which inhibits their applicability (May et al. 1976 and Konigsberger et al. 2015).

In this work, our aim is to overcome such limitations by creating models of metal speciation, including proteins and peptides, and comparing our calculations with metal speciation measurements in simple systems. Our thermodynamic model was created using critically evaluated experimental stability constant data measured over the past 100 years and includes estimates of stability constants made using linear free energy relationships (Prasad & Shock 2025a and Prasad & Shock 2025b). In previous work, we obtained metal speciation in microbial growth media and made correlations with biological response to evaluate metal bioavailability (Prasad & Shock 2025c). Here, we predict metal speciation in blood plasma for several naturally occurring metals and those that may be present as therapeutic agents (metal-based antacid, anti-diabetes and anticancer drugs) or as toxic agents (copper and manganese toxicity). As the medium that transports nutrients and waste products across most human cells, blood plasma is arguably the most crucial biofluid in the human body. Our model replicates the speciation of metals for which elemental distribution in blood plasma is well known and provides metal speciation for cases with extremely limited information on biologically relevant metal-ligand systems. We encourage future experimentalists to test the theoretical predictions made in this work and hope this endeavor furthers research on metal-ligand thermodynamics and metal bioavailability in the human body.

## 2. Calculating speciation

Elemental speciation in any system at equilibrium can be obtained by solving the law of mass-action for all reactions that may happen in the system. So, for the chemical reaction: bB + cC = dD + eE, the law of mass-action is given as:

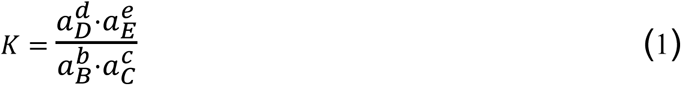

where ‘K’ is the equilibrium constant of the reaction; ‘B’, ‘C’, ‘D’ & ‘E’ are the chemical entities involved in the reaction; ‘a’ is the activity of these entities (also known as ‘corrected concentration’) and ‘b’, ‘c’, ‘d’ & ‘e’ are the stoichiometric constants. As most metal-ligand reactions are complexation reactions, ‘d’ or ‘e’ is 0 and the equilibrium constants are referred to as stability constants. As metal-ligand complexation reactions do not have large kinetic barriers (completing on the order of seconds), the assumption of equilibrium is valid. If the analytical composition (or total composition) of a system is additionally known, the law of mass action can be combined with mass-balance and charge-balance constraints to obtain the chemical distribution of elements occurring in different forms (like calcium as Ca^+2^ or CaOH^+^, CaCl^+^, etc.).

Such speciation calculations can be performed using numerous programs like PHREEQC (Parkhurst 2013) and JESS (May 2015). We have used the software EQ3/6 (Wolery 2010) as it is freely available, and the thermodynamic databases can be easily modified. The composition of blood was obtained from the compilation in the Geigy Tables (Lentner 1981). This composition represents the average blood plasma composition of a healthy human adult and is given in Table 1. As ligand abundance plays a major role in determining metal speciation, we did not include ligands present at concentration < 1µM. The metal-ligand complexes for which experimental equilibrium constants were absent in the literature, were estimated using linear free energy relationships (Prasad & Shock 2025a and Prasad & Shock 2025b). These include metal complexes of albumin, transferrin, glutathione, several amino acids, lactate and ascorbate. A schematic for the flow of information in the procurement of metal-ligand speciation is presented in Fig. 1.

**Table 1.**
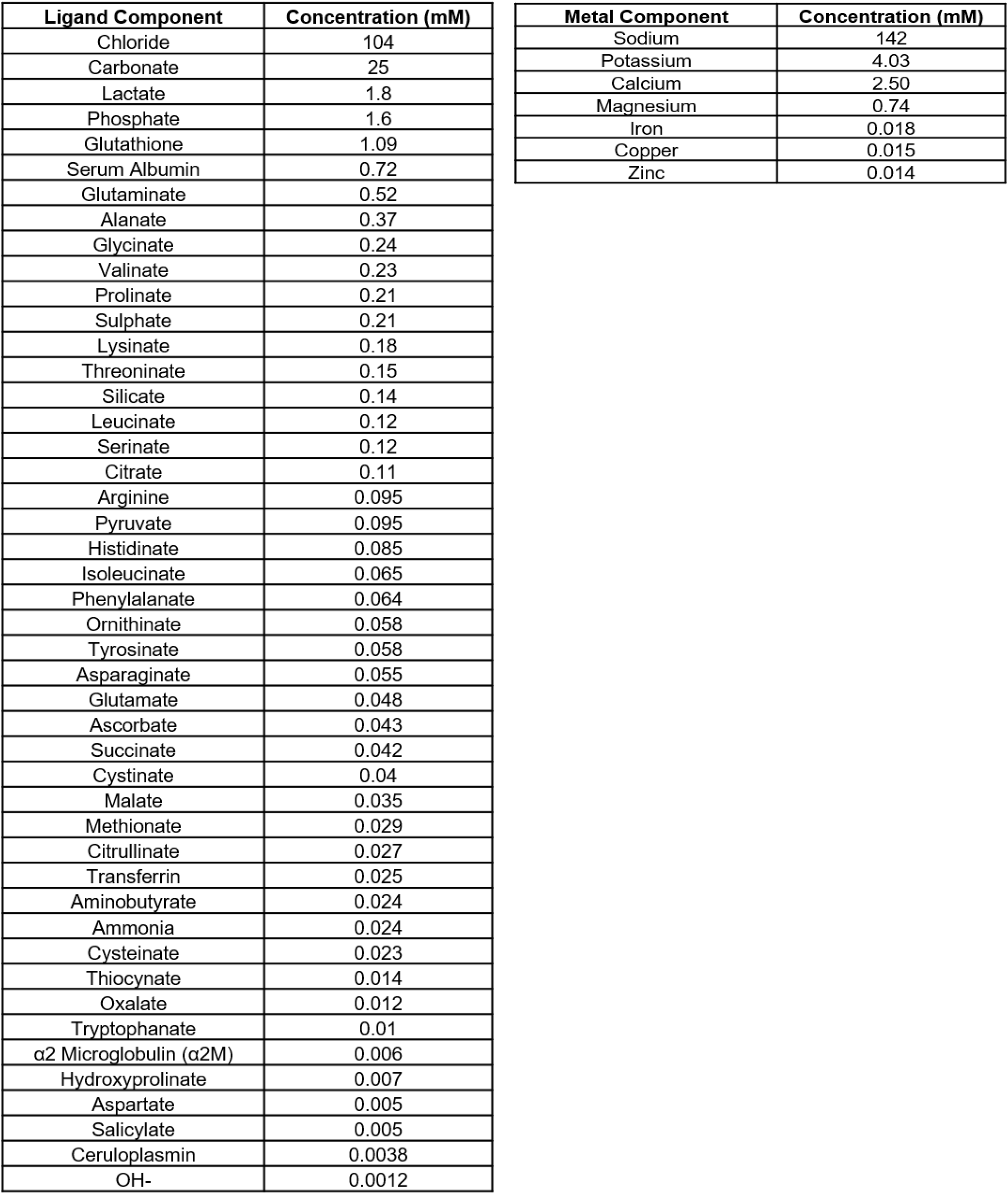
Default composition of blood plasma used in this study.

**Fig 1.**
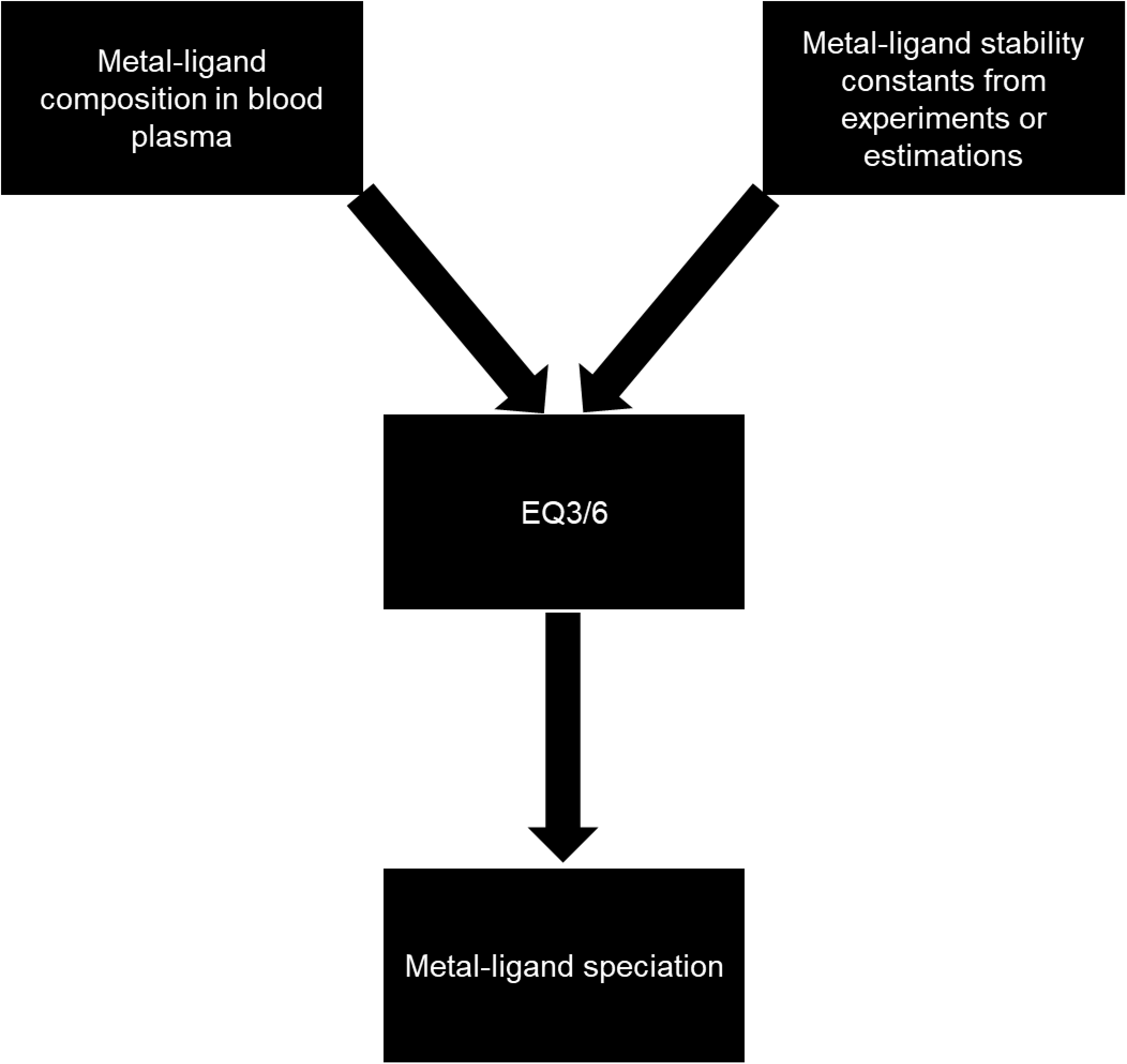
Flowchart for obtaining metal-speciation.

## 3. Speciation of predominant metals in default blood plasma composition

As documented in Table 1, potassium, sodium, calcium and magnesium in an average, healthy human are present at millimolar concentrations while iron, copper and zinc are present at micromolar levels. Owing to their high abundance, the predominant forms of these metals are well established; potassium and sodium are mostly present as free cations, 45% of calcium and magnesium is bound to serum proteins (Peters 1996), almost all of the iron is bound to transferrin (Levina et al. 2017), copper is mostly bound to ceruloplasmin (Hellman & Gitlin 2002) and the majority of zinc is bound to albumin (Kiss et al. 2009). Our speciation calculations reproduce these results and additionally, provide the distribution of these metals in the lesser dominant forms. The speciations for metals present at millimolar and micromolar concentrations are given in Fig. 2a and Fig. 2b, respectively.

**Fig 2(a-b).**
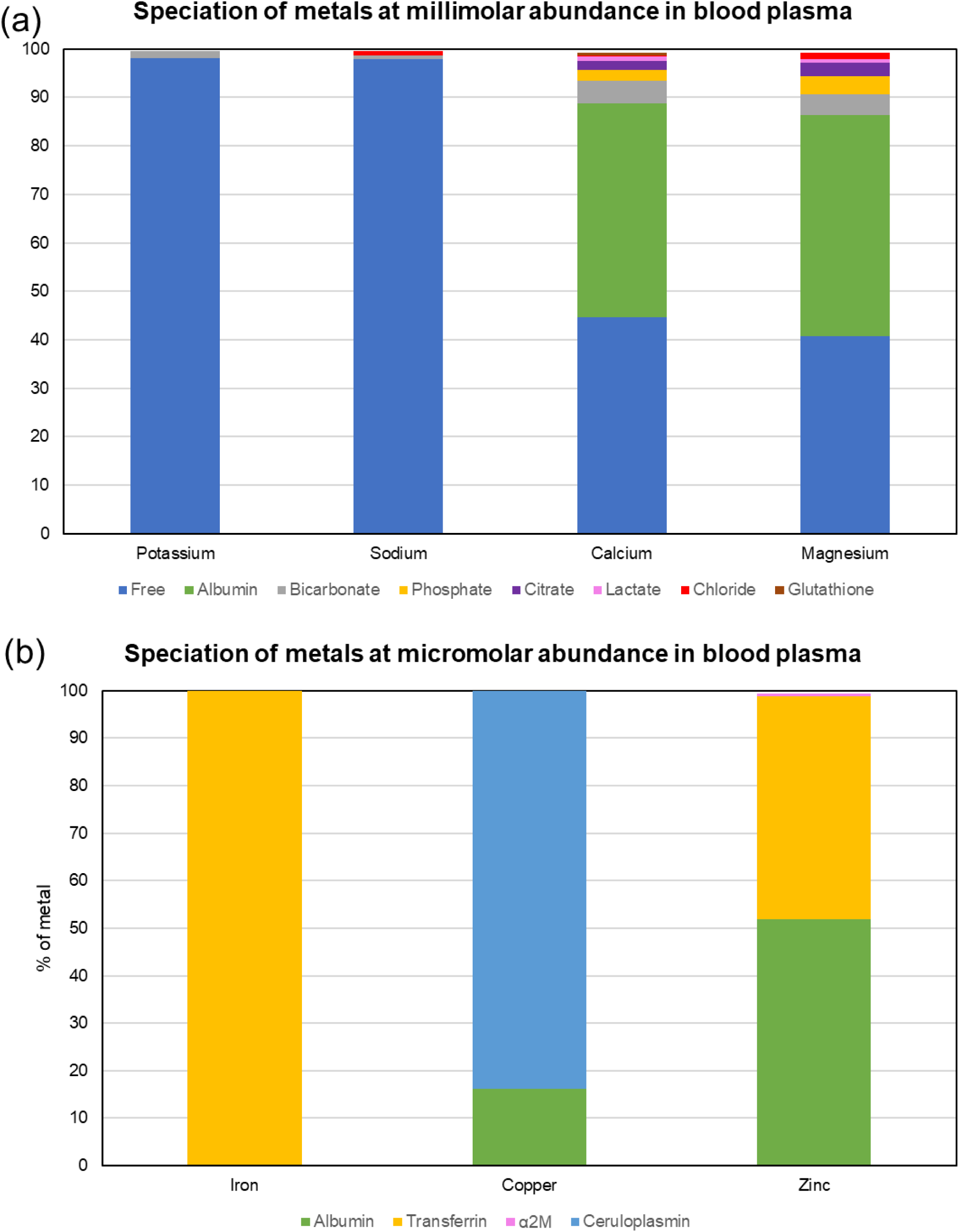
Speciation of metals present at an abundance of (a) 1mM or higher (b) 1µM or higher

Among these seven dominant metals, potassium and sodium are the only ones unaffected by protein complexation. This is understandable due to their low charge and high ionic radii that limit coordination at ‘metal-binding sites’ of proteins. There have been multiple contradictory reports of calcium-albumin stability constants in the literature (McLean & Hastings 1935, Pedersen 1971, Fogh-Andersen 1977 and Kragh-Hansen & Vorum 1993). We found that stability constant values (K=10^2.7^) and number of binding sites (5) reported by Vorum et al. (1995) gave results consistent with the established calcium partitioning in albumin of ∼45 % (Peters 1996).

While the albumin-fraction obtained using the thermodynamic data provided by some studies (Pedersen 1971, Fogh-Andersen 1977 and Kragh-Hansen & Vorum 1993) was slightly lower (by 10-20%), it was extremely low with the data provided by McLean & Hastings 1935 (by ∼40%). Among the remaining calcium, 41% is distributed as the free ion and ∼9% is complexed with bicarbonate, phosphate and citrate, all of which agrees with the overall speciation of calcium (Peters 1996). As magnesium is known to share calcium-binding sites (Peters 1996), we used an identical set of thermodynamic information (same sites and same stability constant values as Ca-albumin) for magnesium-albumin interaction that produced the accepted magnesium distribution in blood plasma (Peters 1996). These results illustrate how speciation calculations may be used to critically evaluate ambiguous or limited thermodynamic data.

As can be seen from Fig. 2(b), transferrin dominates iron speciation in blood plasma. Iron(III)-transferrin interaction has been well-studied and the stoichiometric constants (reaction quotient expressed with concentration) reported by Aisen et al. (1978) are frequently used in the literature. However, these values vary with pH and do not qualify as thermodynamic equilibrium constants that are defined as reaction quotient expressed with activity. Additionally, the investigation precluded acid association constants (pK_a_) of amino acids at the metal-binding site. Subsequent research (Sun et al. 2004) shows that the pK_a_ of one tyrosine at the site (Tyr188) is considerably lower (7.2) than the pK_a_ of the free tyrosine sidechain (10.2). The pK_a_ of the histidine at the site (His249) was found to be comparable to imidazole pK_a_ (Woodworth et al. 1987, Valcour & Woodworth 1987 and Kubal et al. 1994) and those of the other amino acids (Tyr95 and Asp63) were assumed to be same as the corresponding sidechain pK_a_. As transferrin has two metal-binding sites, the same considerations were made for the other site (Tyr517, His585, Tyr426 and Asp392). These pK_a_ values were incorporated in our database along with the corresponding thermodynamic data associated with the competitive-ligand citrate which allowed us to obtain thermodynamic stability constant values for Fe(III)-transferrin by regression of experimental data reported by Aisen et al. (1978) (see Appendix). While the use of these thermodynamic stability constants does not change the iron speciation at the pH of blood plasma (Fig. 2b), it has substantial effects on iron speciation at lower pH (see below).

While most investigations into copper speciation agree that the protein ceruloplasmin dominates copper speciation in blood plasma, the fraction of copper as ceruloplasmin ranges from 65% (Linder et al. 1996) to 95% (Hellman & Gitlin 2002). Additionally, the thermodynamic information for metal-ceruloplasmin complexes is very limited. While it is well agreed upon that ceruloplasmin has six primary sites for copper (Hellman & Gitlin 2002), data on stability constants are obscure at best. Speciation calculations with data reported by Zgirski & Frieden (1990) underestimate copper-ceruloplasmin abundance by several orders of magnitude. Subsequent X-ray crystallography investigations revealed a preponderance of cysteine and histidine groups in the three ‘types’ of copper sites in the protein (Hellman & Gitlin 2002) suggesting that stability constant values are much larger than the reported values (K∼10^7^). Additionally, as ceruloplasmin dominates copper speciation despite the higher concentration of serum albumin, Cu-ceruloplasmin stability constants should be larger than Cu-albumin stability constants that are known to be ∼ 10^12^-10^13^ for the N-terminus site (NTS). We adopted an average stability constant of 10^14^ for the six Cu-ceruloplasmin sites that results in the speciation shown in Fig. 2(b). Cu-ceruloplasmin constitutes ∼84% of the total copper while ∼15% is distributed as Cu-albumin. Copper speciation is extremely sensitive to this stability constant value as chancing K from 10^13.5^ to 10^14.5^ resulted in Cu-ceruloplasmin varying across the established speciation range of 65% to 95%.

Through modeling calculations, it is largely established that almost all zinc in blood plasma (∼98%) is bound to the serum proteins albumin, α2-microglobulin (α2M) and transferrin (Kiss et al. 2017). However, the ratio of zinc distribution among these proteins is still ambiguous. To resolve this issue, we simulated zinc speciation measurements associated with these proteins using stability constants reported in the literature while obtaining stability constants by regression. Zn-albumin stability constants reported by Lu et al. (2012) for the Multi-Binding Site and for Site B by Ohyoshi et al. (1999) agree closely with the experimental measurements reported by Bytzek et al. (2009) (see Appendix). Zn-transferrin stability constants were obtained using a procedure similar to the method used to obtain values for Fe(III)- transferrin based on experimental measurements reported by Harris & Stenback (1988) (see Appendix), while stability constants reported by Adham et al. (1977) were used for Zn-α2M complexes. Using the above set of data and the stability constants for other ligands mentioned previously, we predict that ∼95% of zinc in blood plasma is bound to serum proteins (Fig. 2b). While this agrees with the value of Zn-protein fraction established in the literature (Bytzek et al. 2009, Lu et al. 2012 and Kiss et al. 2017), we found that transferrin takes up ∼40% of the total zinc, which is much higher than the conventional values (Bytzek et al. 2009). Most papers in the literature report that albumin takes up 70-85% of zinc while α2M constitutes ∼10-30% of total zinc (Falchuk et al. 1977, Charlswood 1979, Kiilerich & Christiansen 1986, Boyett & Sullivan 1970, and Foote & Delves 1984). However, these studies were performed before the experiments of Harris & Stenback (1988), which was the first study that carefully evaluated Zn-transferrin association. Additionally, none of these studies evaluated zinc distribution in the presence of all three proteins. Moreover, while the stability constants for Zn-transferrin and Zn-α2M are comparable, transferrin is more abundant than α2M in blood plasma by a factor of 6 (Table 1). For these reasons, we predict that our calculated speciation is closer to the *in vivo* zinc speciation and encourage future researchers to investigate this subject via experimentation. Our fraction of Zn-albumin (∼53%) is closer to the values reported by Peters (1996) and Vallee & Falchuk (1993) (65-70%).

While the metals mentioned above are evidently needed, major changes in their abundance and distribution can severely affect human health either profitably or adversely. In the subsequent sections, we describe several cases where metals are investigated for their therapeutic potential or their toxicity. We have predicted the speciations of these metal-based drugs and explored their dependence on pH, ligand concentration, and metal abundance.

## 4. Metal speciation in therapeutic or toxic scenarios

### 4.1 Gallium-based anticancer drugs

Gallium-based compounds have been a topic of interest in anticancer research for almost 55 years (Hart & Adamson 1971, Foster et al. 1986, Chitambar & Sax 1992, Chen et al. 2007 and Cao et al. 2019). Besides being agents of cytotoxicity, they are also used in cancer research for tumor-imaging owing to the radioactive properties of isotopes ^67^Ga and ^68^Ga (Edwards & Hayes 1969, Jackson & Byrne 1996, Schuhmacher et al. 2001, McInnes et al. 2017 and Yaxley et al. 2019). While it is generally accepted that lipophilic complexes of gallium increase its cellular uptake in comparison to inorganic salts (Kiss et al. 2017 and Levina et al. 2017), there is limited work establishing the dependence of gallium bioavailability on its speciation and dosage in bodily fluids. Speciation is of direct relevance because metal-based anticancer drugs are known to produce long-lasting anticancer immune responses (Englinger et al. 2019). Here, we report predicted gallium speciation in blood upon addition as Ga(Maltol)_3_ and compare our calculations with experimental results and models from the literature.

The antineoplastic effects of Ga(Maltol)_3_ were first reported in 2000 (Bernstein et al. 2000) and even reached Phase I-II of clinical trials in United States in 2005 (https://clinicaltrials.gov/ct2/show/study/NCT00050687). Even though the results of the clinical trials have not been published, subsequent research on Ga(Maltol)_3_ cytotoxicity has generally produced favorable results (Chua et al. 2006, Chitambar et al. 2007, Merli et al. 2018 and Chitambar et al. 2018). Perhaps one of the reasons for the continued interest in Ga(Maltol)_3_ research is its high oral bioavailability. Due to its relatively high lipophilicity (neutral overall charge and aromatic chelate rings), it has the tendency to cross the lipid barrier via passive diffusion. This phenomenon seems to be ligand dependent as maltol complexes of aluminum and iron are also known to have high bioavailability (Finnegan et al. 1986, Finnegan et al. 1987, Reffitt et al. 2000, Murukami et al. 2006, Pereira et al. 2014 and Stallmach & Buning 2015). The mechanism of action of Ga(III)-based anticancer drugs has largely been attributed to its Fe(III)- like properties. As cancer cells have a high iron dependence, this leads to the disruption of several iron-dependent processes in the cell like iron homeostasis, iron-dependent ribonucleotide reductase activity and the activation of Bax protein that triggers apoptosis through the mitochondrial release of cytochrome c and caspase-3 (McInnes et al. 2017, Chitambar 2017 and Englinger et al. 2019). For these reasons, Ga(Maltol)_3_ is still a sought-after drug in anticancer research.

Despite the large body of research on Ga(Maltol)_3_, we found only one study that investigated its speciation. Enyedy et al. (2015) evaluated the interaction between Ga(Maltol)_3_ and the two predominant serum proteins albumin and transferrin using multiple techniques. They also reported conditional thermodynamic constants for drug-protein binding (Kiss et al. 2009) and computed gallium distribution as a function of Ga(Maltol)_3_ concentration (Enyedy et al. 2015). However, this distribution was reported for a relatively simple system comprising of only four ligands (maltol, albumin, transferrin and hydroxide). To investigate if other common ligands in blood like citrate, lactate and amino acids affect gallium distribution, we performed gallium speciation calculations for the default composition of blood reported in Table 1. We found that our results for gallium distribution closely mirrored the speciation reported by Enyedy et al. (2015) as illustrated in Fig. 3a. This suggests that the LMM (low molecular mass) ligands do not significantly impact gallium speciation at the physiological pH of 7.4. However, we found that gallium speciation was completely different at lower pH as shown in Fig. 3b. For example, for 100µM Ga(Maltol)_3_ at a pH of 4, which is close to the pH of endosomes, only about 45% of gallium is bound to transferrin whereas Ga(Maltol)_3_^0^ takes up only ∼9% gallium. The rest of the gallium is predominantly distributed as Ga(Maltol)_2_^+^ and citrate complexes that may have a different biological fate in comparison to the lipophilic Ga(Maltol)_3_^0^ and the transferrin-bound gallium. At pH>7.4, gallium is predominantly distributed as Ga(OH)_4_^-^, which is consistent with results found in the literature (Harris & Pecoraro 1983).

**Fig 3(a-b).**
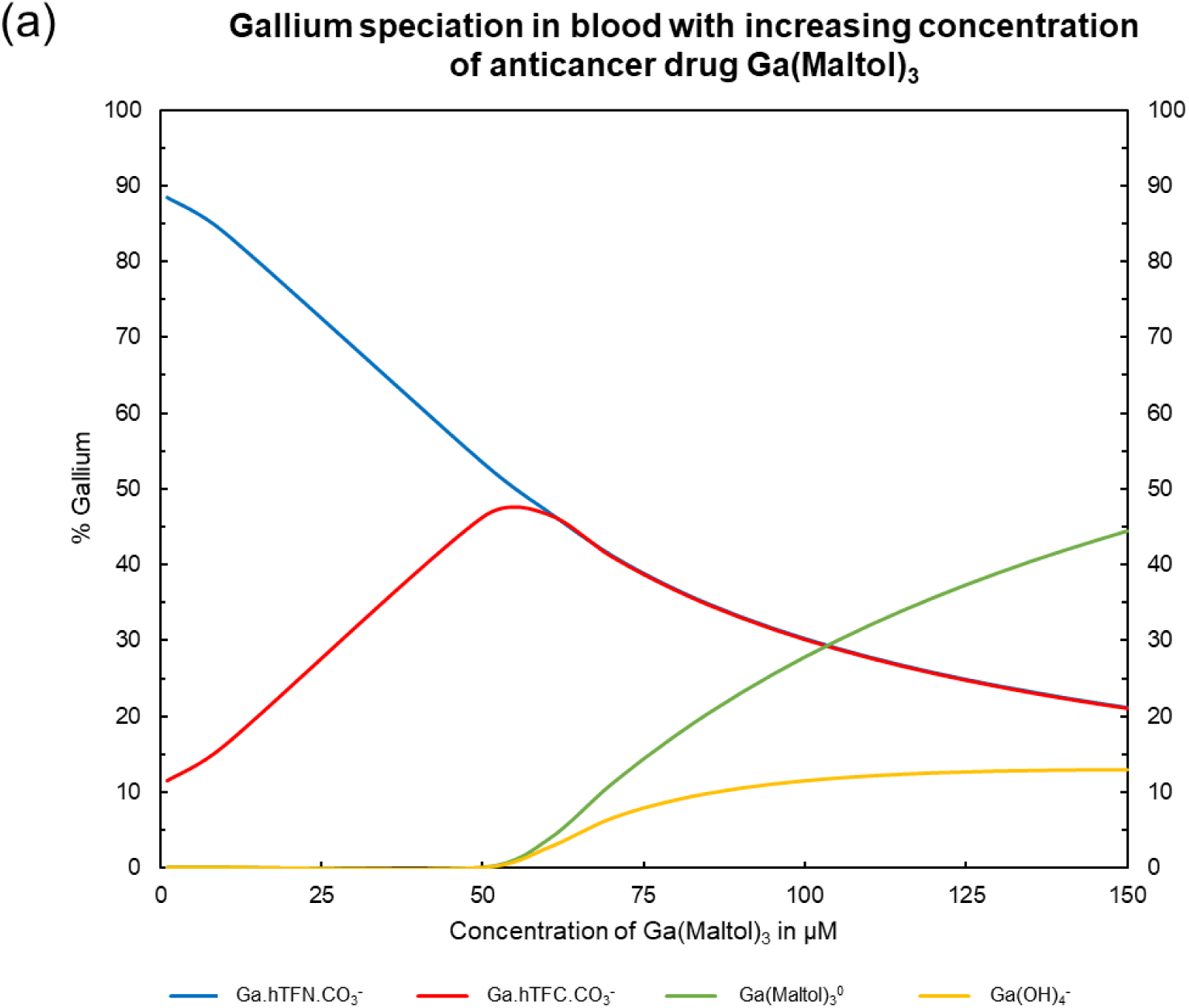
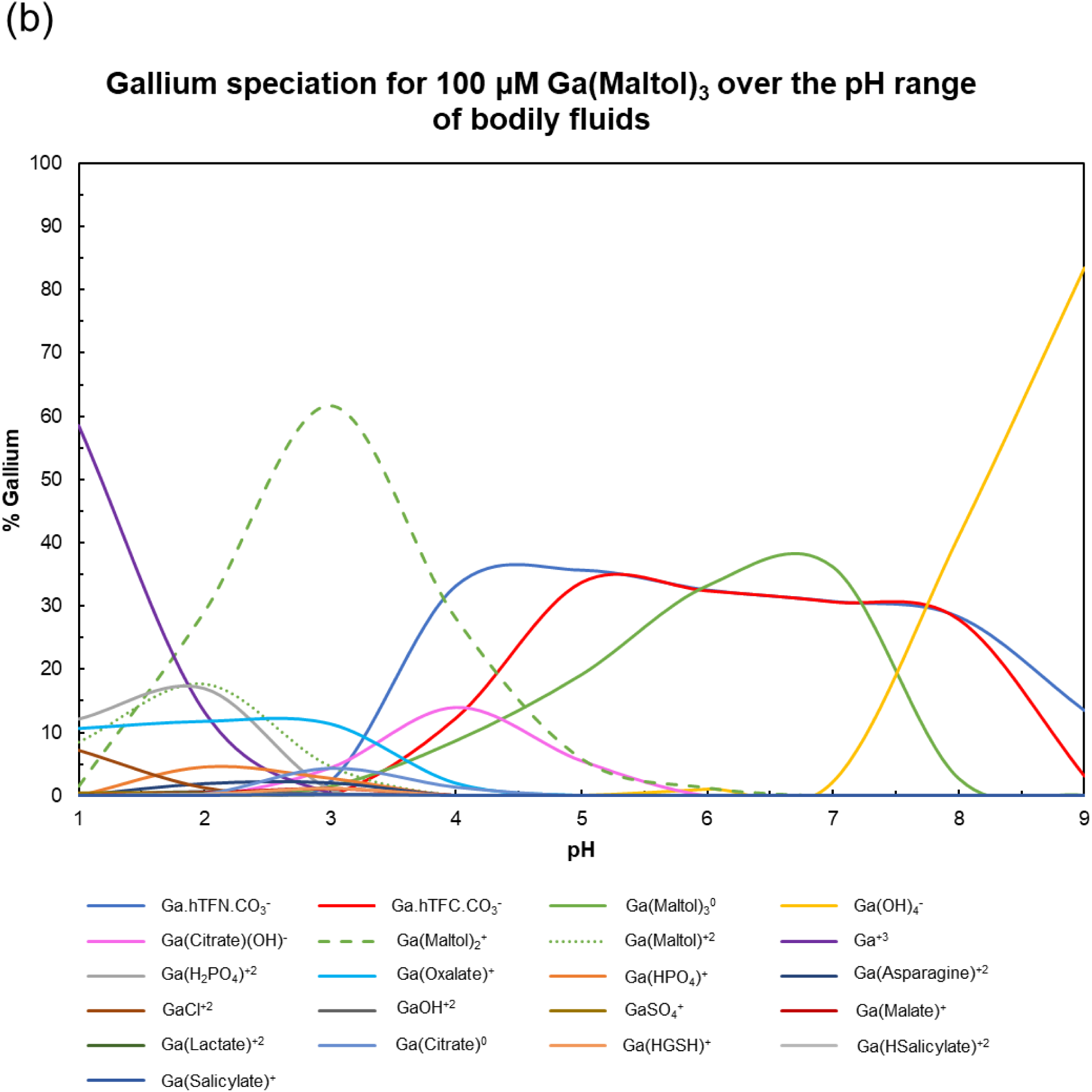
Speciation and bioavailability of gallium-based anticancer drugs

The results discussed above show how thermodynamic models of metal-based drugs can provide useful information on their distribution and bioavailability in the human body. Such models can also be made for the next generation of gallium-based anticancer drugs currently under preclinical trial like corroles, azoles and thiolates (Chitambar 2017). As speciation calculations are rapid and inexpensive, they are an excellent means to selecting metal-based drugs with high bioavailability. Experimental research performed in conjunction with these calculations can constrain dosage levels for these anticancer drugs that elicit cytotoxic tumor response without causing long-lasting anticancer immune responses.

### 4.2 Copper dysregulation and Wilson Disease

Owing to its redox nature and high crustal abundance, all living cells use copper as a source and acceptor of electrons at the active sites of many enzymes like cytochrome c oxidase and superoxide dismutase. Therefore, copper is an essential trace metal, vitally important for human health and well-being. Copper deficiency (hypocupremia) in humans is known to cause myelopathy that is known to be associated with Vitamin B_12_ deficiency (Kumar et al. 2004, Prodan et al. 2007) while excess of copper (hypercupremia) may lead to cancer (Buchwald & Hudson 1944, Margalioth et al. 1983, Gupte & Mumper 2009). The plasma copper concentration associated with these diseases ranges from 1.5µM in hypocupremia to 50 µM in hypercupremia (Lentner 1981). Our predicted speciation for copper in blood plasma under these conditions is shown in Fig. 4a. Note that most of the copper is distributed between the proteins ceruloplasmin and albumin (at the NTS site). As the concentration of albumin (0.63mM) is much higher than that of ceruloplasmin (3µM), the Cu-albumin fraction increases as total copper in blood plasma increases. As the established role of the NTS on albumin is to transfer copper from the intestine to liver (Bal et al. 2013), this may have implications on hepatotoxicity. As albumin is regarded as a primary component of the exchangeable copper in the plasma (Linder & Hazegh-Azam 1996), the high Cu-albumin fraction could explain why copper is toxic despite being completely distributed among plasma proteins.

**Fig 4(a-c).**
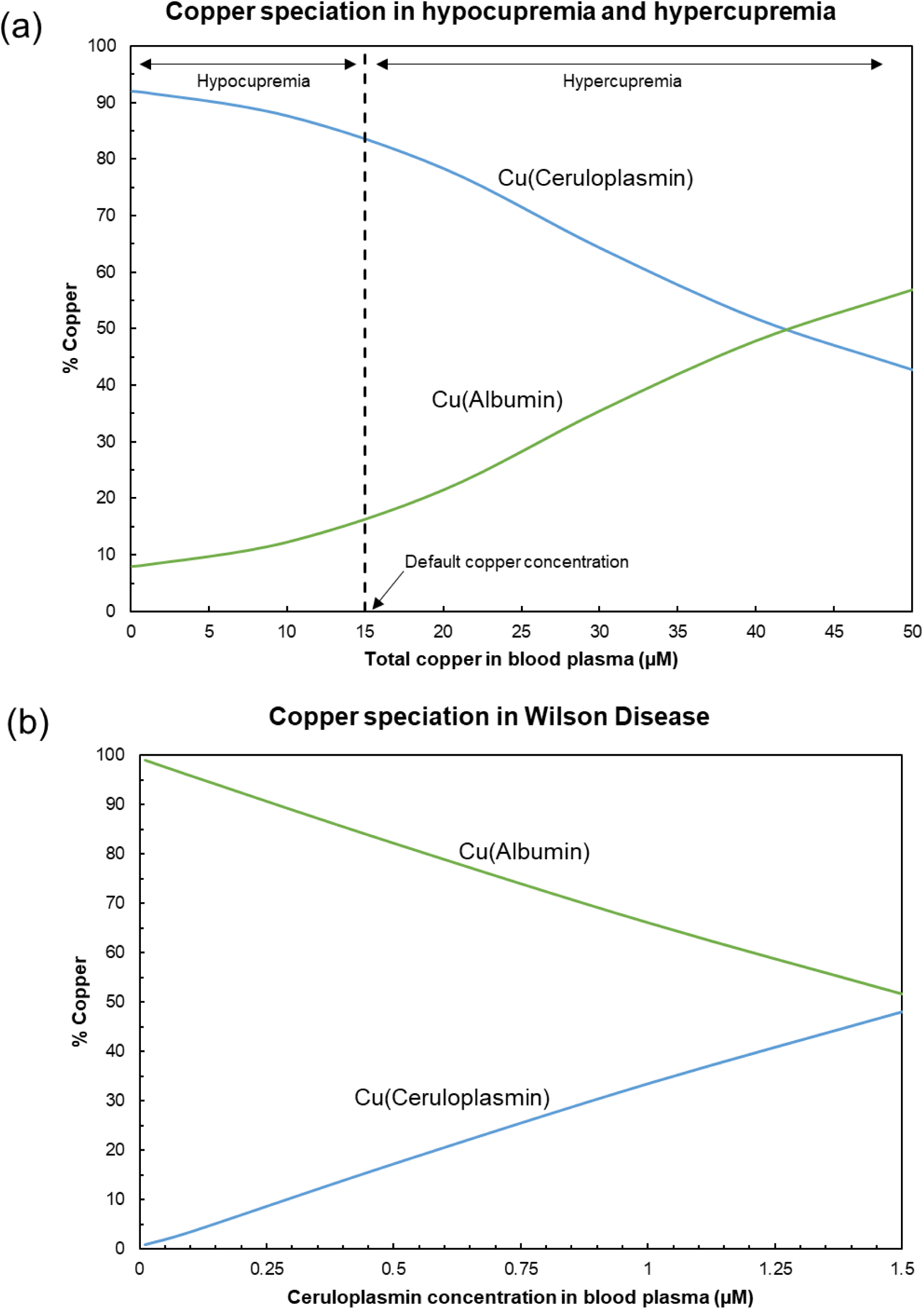
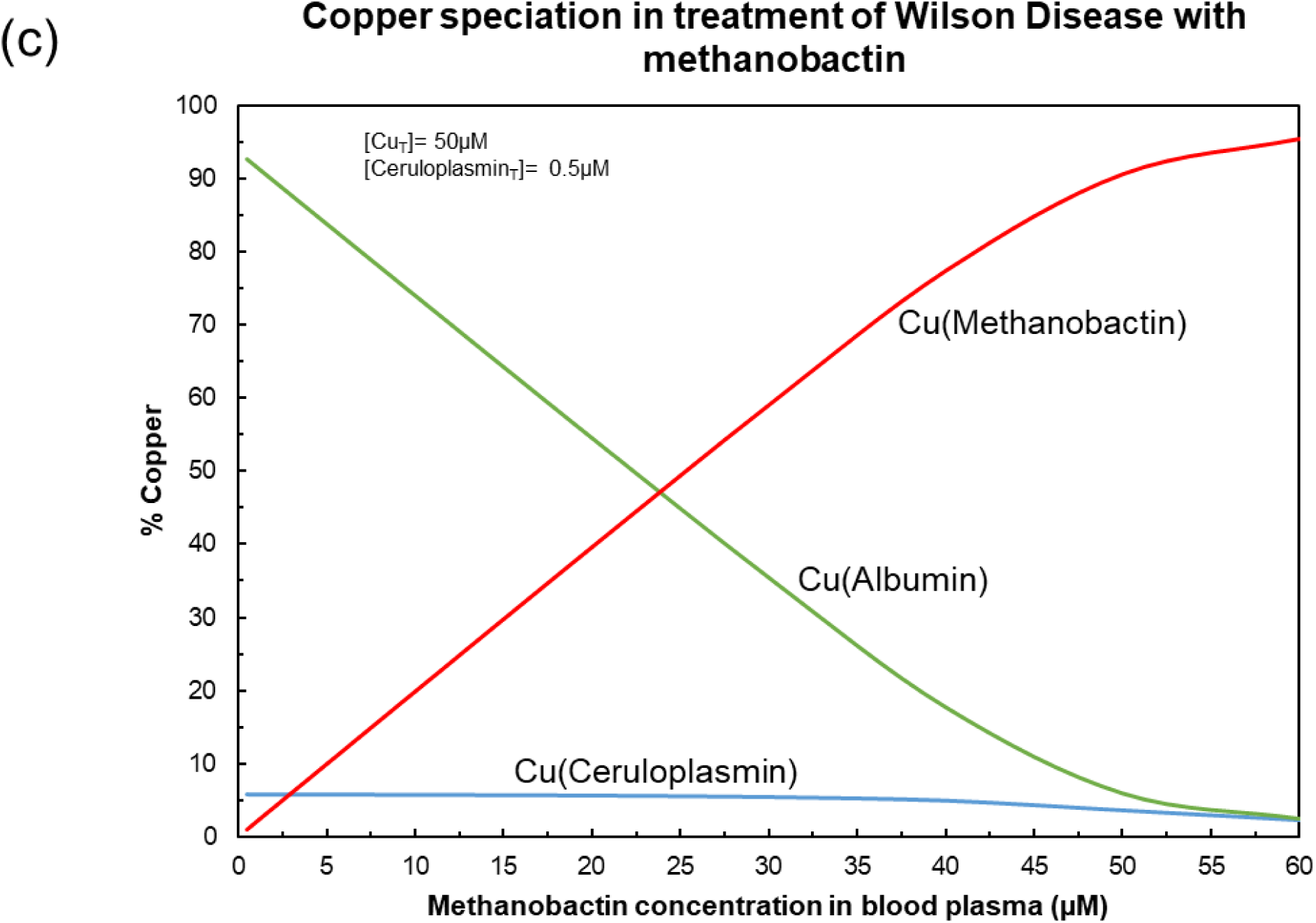
Copper(II) speciation in blood plasma under different conditions.

Copper dysregulation in humans may also occur due to genetic reasons. Mutations in the gene encoding for the ATP7A protein causes Menkes Disease that leads to impaired copper absorption in the blood (Prohaska 2008). On the other hand, Wilson Disease (or Wilson’s Disease) is caused by disabling mutations in both copies of the gene encoding for the ATP7B protein, which leads to copper excess mainly in the liver and brain (Bull et al. 1993, Prohaska 2008). As Wilson Disease is about ten times more prevalent than Menkes Disease (de Bie et al. 2007), we have focused our attention on Wilson Disease in this work.

One of the key characteristics of Wilson Disease (WD) is deficiency of ceruloplasmin in blood plasma (Scheinberg & Gitlin 1952, Hellman & Gitlin 2002). The low ceruloplasmin concentration in patients suffering with WD is a consequence of the lack of functional ATP7B in hepatocytes, resulting in the secretion of a rapidly degraded apoprotein (Gitlin 2003). Generally, individuals suffering from WD have ceruloplasmin concentration lower than 1.3µM, which is about 30% of normal levels (Gitlin 2003). The predicted dependence of plasma ceruloplasmin abundance on copper speciation is demonstrated in Fig. 4b. Note that throughout this range of ceruloplasmin concentration, the Cu-albumin fraction dominates over the Cu-ceruloplasmin fraction, which may explain the copper toxicity associated with the disease.

There are primarily two treatments for Wilson Disease-chelation therapy and zinc supplementation. For patients who are unresponsive to both treatments, liver transplantation is the only recourse. Zinc supplementation inhibits the absorption of copper in the gastrointestinal mucosa and stimulates metallothionein synthesis in the enterocytes that have a higher capacity to chelate dietary copper (Patil et al. 2013, Delangle & Mitz 2012, Brewer 2009). While this strategy has limited side-effects, its relative slow response makes it a suitable method of treatment at the presymptomatic stage (Huster 2009, Linn et al. 2009). Chelation therapy is the preferred form of treatment in the clinical stages of Wilson Disease. Several copper-chelators have been investigated as putative WD drugs such as penicillamine, triethylenetetramine and tetrathiomolybdate (Delangle & Mitz 2009). Recently, the naturally produced chalkophore methanobactin was used as a chelator to reverse liver failure in Wilson Disease (Lichtmannegger et al. 2016, Summer et al. 2011). Methanobactin (Mb) is a 1154 Da post-translationally modified peptide produced by the methanotroph *Myethylosinus trichosporium* OB3b to accumulate copper that is essential for the activity of the methane monooxygenase enzymes (Morton et al. 2000, Knapp et al. 2007, Semrau et al. 2010, Kenney & Rosenzweig 2013). It chelates copper with a 1:1 stoichiometry with an extremely large stability constant (K= 10^20.8^, El Ghazouani et al. 2011). While other metals like cobalt, nickel and zinc can also be chelated by Mb (Choi et al. 2006), the nitrogen-sulfur N_2_S_2_ coordination imparts a strong selectivity for copper consistent with its Cu detoxifying ability. The predicted effect of methanobactin on copper speciation in an extreme case of WD is illustrated in Fig. 4c. The values for methanobactin protonation constants (pK_a_) were taken from Pesch et al. (2012). Using a total plasma copper concentration of 50 µM and total plasma ceruloplasmin concentration of 0.5 µM, our model predicts that ∼ 60 µM (∼0.07g) methanobactin can restore the exchangeable copper abundance on albumin to normal levels. Similar models can be made for specific cases of copper and ceruloplasmin abundance in plasma of WD patients. Evidently, similar models need to be made for bile, intracellular hepatocyte composition and urine to obtain a deeper understanding of copper distribution in this disease. We hope these results encourage future researchers to conduct further experimental investigations on this topic in conjunction with models of copper speciation.

### 4.3 Zinc-based antidiabetics

The role of zinc in diabetes is a topic of considerable interest in the current scientific community with several recent studies exploring the associated physiology, molecular machinery and genetics (including: Nazem et al. 2019, Bosma et al. 2019, Jonsdottir et al. 2019, Adulcikas et al. 2019, Sobczak et al. 2019, Patel et al. 2019). While many researchers agree that zinc supplementation has a beneficial effect on diabetes patients (Wang et al. 2019, Nazem et al. 2019), others are more skeptical of the benefits of additional zinc intake (Fernandez-Cao et al. 2019, Perez et al. 2018). Some reports suggest that diabetic physiochemistry is associated with dysregulation of zinc homeostasis rather than total zinc levels (Fukunaka & Fujitani 2018, Chu et al. 2017, Fernandez-Cao et al. 2018). Thus, although the link between zinc and diabetes is broadly recognized, a deeper understanding may emerge through investigation of zinc speciation.

Some studies have obtained zinc speciation with multiple putative carrier-ligands that might increase zinc-bioavailability and hence elicit high anti-diabetic activity (Enyedy et al. 2008, 2012, Kiss et al. 2009, 2012, Bytzek et al. 2009). However, these studies were performed in simple aqueous solutions containing of only 5-6 ligands. We realized that obtaining zinc speciation in a composition more reflective of blood plasma would enhance the current state of knowledge for prospective zinc-based antidiabetic drugs. Therefore, we predicted zinc-speciation in blood plasma including the ligand oxine, a commonly used carrier-ligand.

Zn-oxine stability constants were taken from Nasanen & Penttinen 1952 and Fresco & Freiser (1964), while the oxine-albumin stability constant was taken from Enyedy et al. 2015. Stability constants for other complexes were taken from the literature or estimated using linear free energy relationships, as summarized by Prasad & Shock 2025b. These stability constants along with the composition described in Table 1 were used to obtain the speciation of zinc as a function of the total Zn(Oxine)_2_ in blood plasma shown in Fig. 5a. As can be seen in the figure, the abundance of the neutral aqueous complex Zn(Oxine)_2_^0^ changes in a non-linear manner with increasing concentration of total Zn(Oxine)_2_ in blood plasma. Additionally, note that Zn(Oxine)_2_^0^ dominates zinc speciation above 450µM, which is quite high. Adjudicating a suitable dose for this prospective drug is enabled by investigating the bioavailability of zinc in solutions of Zn(Oxine)_2_.

**Fig 5(a-b).**
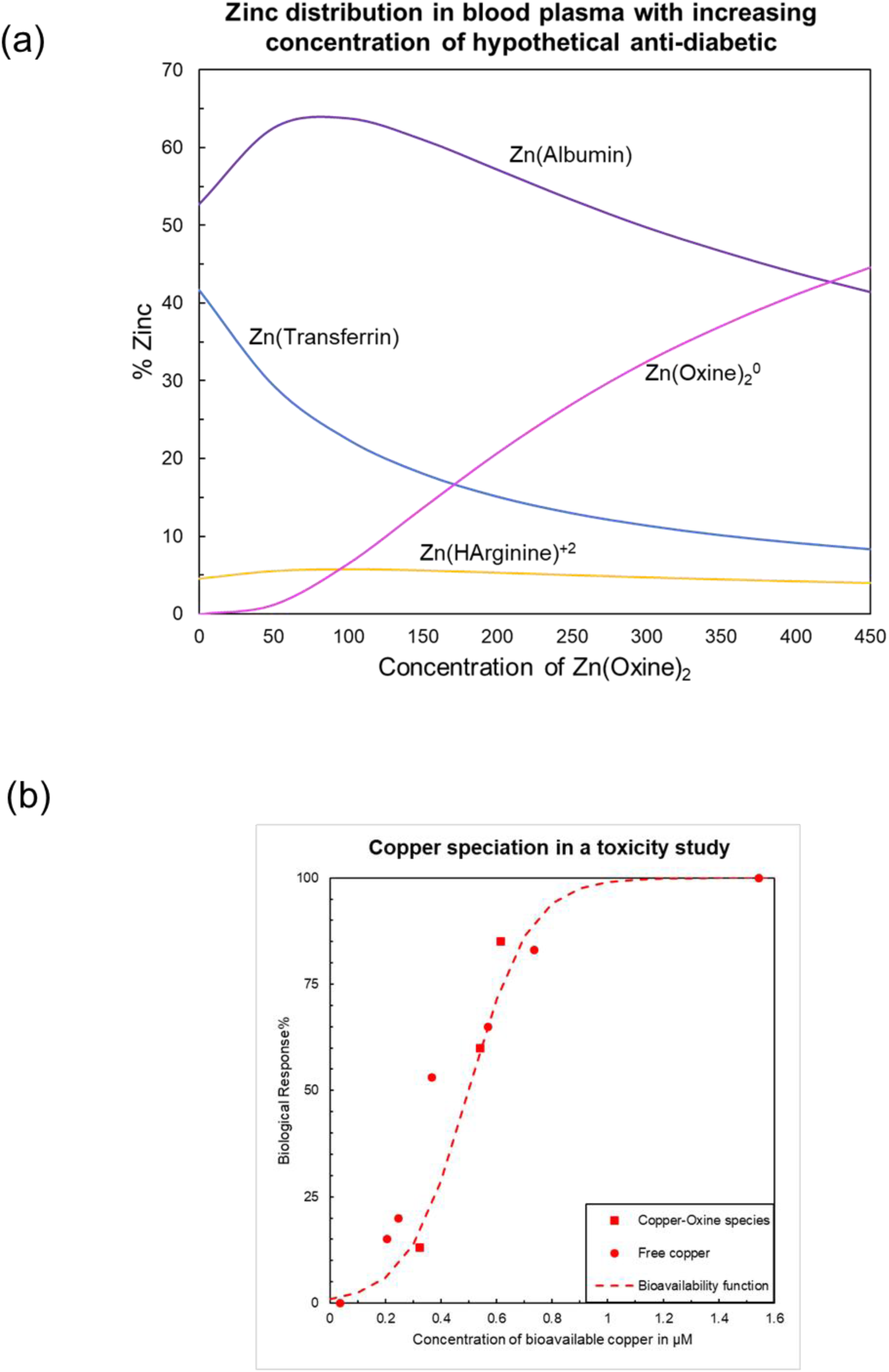
Speciation and bioavailability of zinc-based antidiabetics.

The bioavailability of many metal-based drugs depend on the form in which they are administered. For example, iron administered as a maltol complex is known to be highly bioavailable in human cells (Stallmach & Buning 2015, Pereira et al. 2014, Murukami et al. 2006, Reffitt et al. 2000). Choosing ligands like oxine and maltol as carrier-ligands enhances the lipophilic/hydrophobic nature of the molecules. As an example, Zn(Oxine)_2_^0^ consists of a polar nucleus consisting of the zinc ion and polar functional groups of the two oxine molecules (oxygen and nitrogen). The outer regions of this binary complex are two aromatic rings that have high lipophilicity. This type of structure allows such molecules to pass though the hydrophobic lipid bilayer via passive diffusion. This phenomenon was demonstrated in an amphipod copper toxicity study (Ahsanullah & Florence 1984), for which we obtained the bioavailability curve Fig. 5b consisting of free copper and copper-oxine complexes (Prasad & Shock 2025c). As can be seen from the figure, the biological response was extremely low when the concentration of the bioavailable metal was 0-0.3 µM but increased considerably thereafter. Given the similar behavior of Cu^+2^ and Zn^+2^, we propose that a similar response can be expected with the administration of zinc-oxine complexes.

One way of determining the anti-diabetic activity of a drug is by measuring the inhibition of free fatty acid (FFA) release in adipocytes upon addition of epinephrine (Nakai et al. 1995). This property was measured by Sakurai et al. 2002 for a variety of metal ions including Zn^+2^ in the presence of insulin from mice that had been administered zinc-based compounds (3mg per kg of mouse weight) for two weeks. The inhibition of free fatty acid release for Zn^+2^ was measured to be ∼23%, that was only slightly lower than the corresponding values for V^+3^ and VO^+2^ compounds that have been antidiabetic candidates for several decades. The zinc-based compounds investigated included Zn(Maltol)_2_, Zn(Picolinate)_2_ and Zn(6-methyl picolinate)_2_, all of which can pass through the lipid bilayer via passive diffusion like Zn(Oxine)_2_. Given that the lipid bilayer is universal to life, it may be postulated that bioavailability of lipophilic complexes will be similar in amphipods and human cells. Combining this assumption with the information provided by Sakurai et al. 2002, we obtain that 60 µM of total Zn(Oxine)_2_ in blood plasma is enough to elicit the maximum possible anti-diabetic zinc activity as it is present at > 1 µM abundance., This dosage is more moderate in comparison to the recommended dose of Nazem et al. 2019. Arriving at a suitable dosage could be attained by combining experiments and speciation calculations across the digestive system to account for the change in Zinc(Oxine)_2_ speciation from the concentration in mouth to the concentration in blood.

### 4.4 Bismuth-based antacids

For over 200 years, bismuth-based compounds have been used worldwide to treat gastrointestinal complaints like dyspepsia and diarrhea (Chambers 1875, Bierer 1990). However, bismuth-based drugs are also known to cause nephrotoxicity with earliest reports dating back to 1802 (Leonard 1926, Gryboski & Gotoff 1961, Czerwinski & Ginn 1964, Cengiz et al. 2005, Liu et al. 2018). Additionally, bismuth compounds are known to cause neurotoxicity and brain encephalopathy as demonstrated by the bismuth epidemic of the 1970s that killed over 100 people in France and Australia (Morrow 1973, Burns et al. 1974, Martin-Bouyer et al. 1981). Despite its toxic history, bismuth compounds continue to attract attention for therapeutic purposes such as eradication of the stomach-infesting bacteria *Heliobacter pylori* (Gisbert 2011, Fallone et al. 2016, Gu et al. 2019), as broad-spectrum inhibitors of antibiotic resistant enzymes like metallo-β-lactamase (Wang et al. 2018), as non-steroidal anti-inflammatory drugs (Hawksworth et al. 2014), and as anticancer agents (Sathekge et al. 2017, Cheng et al. 2018, Ahamed et al. 2019, Chan et al. 2019). Thus, owing to the dual nature of bismuth, regulating its dosage should help to maximize the beneficial effects while minimizing the toxic side-effects.

While the bioavailability of bismuth-based compounds has been discussed for many years, limited research exists on bismuth speciation in human bodily fluids (Bierer 1990, Tillman et al. 1996, Michalke et al. 2009). We found only one study in which the investigators demonstrated that transferrin controls bismuth speciation in blood plasma (Montavon et al. 2012). We found no results on bismuth speciation in cerebrospinal fluid (CSF) despite its direct implications to neurotoxicity and brain encephalopathy. To bridge this gap, we predicted bismuth speciation in the two bodily fluids and the endosome that transfers bismuth across the blood-brain barrier via receptor-mediated endocytosis. Implications on bismuth bioavailability and neurotoxicity are discussed below.

As six recent studies reported cases of bismuth-induced encephalopathy due to the intake of the commercially available form of bismuth subsalicylate (Reynolds et al. 2012, Masannat et al. 2013, Siram et al. 2017, Ali et al. 2017, Hogan et al. 2018, Borbinha et al. 2019), we performed speciation calculations reflecting increasing concentrations of this drug. Bismuth-transferrin stability constants were obtained by regressing stoichiometric constants reported by Li et al. 1996 in conjunction with subsequently measured transferrin pK_a_ values (Appendix). Since no measurements for bismuth-salicylate stability constants exist in the literature, these values were estimated using linear free energy relationships reported by Prasad & Shock 2025b and Shock & Koretsky 1995. Stability constants for other metal-ligand complexes were obtained from compilation or estimates reported in Prasad & Shock 2025b. It should be noted that for most ligands present in blood plasma, stability constants for bismuth complexes were estimated due to the scarcity of experimental measurements.

Predicted speciation of bismuth from nanomolar to millimolar concentration in blood and cerebrospinal fluid are shown in Fig. 6a and 6b, respectively. Comparison of these plots shows that bismuth speciation is completely different in the two bodily fluids-while transferrin dominates the speciation in blood, Bi(OH)_3_ is the predominant form of bismuth in cerebrospinal fluid. A major contributing factor is the substantially lower concentration of transferrin in cerebrospinal fluid (∼0.17 µM) compared to the corresponding concentration in blood (∼ 37 µM). As neutral LMM complexes tend to pass through the lipid bilayer via passive diffusion (Levina et al. 2017), Bi(OH)_3_ may be more readily bioavailable than other forms of bismuth. Recently, Bi(OH)_3_ nanoparticles were found to possess high toxicity against malignant cancer cells (Bogusz et al. 2018). It is therefore likely that the corresponding aqueous form is also cytotoxic, which may also help explain the neurological dysfunction associated with this metal. We suggest that additional studies of the influence of Bi(OH)_3_ are warranted.

**Fig 6(a-f).**
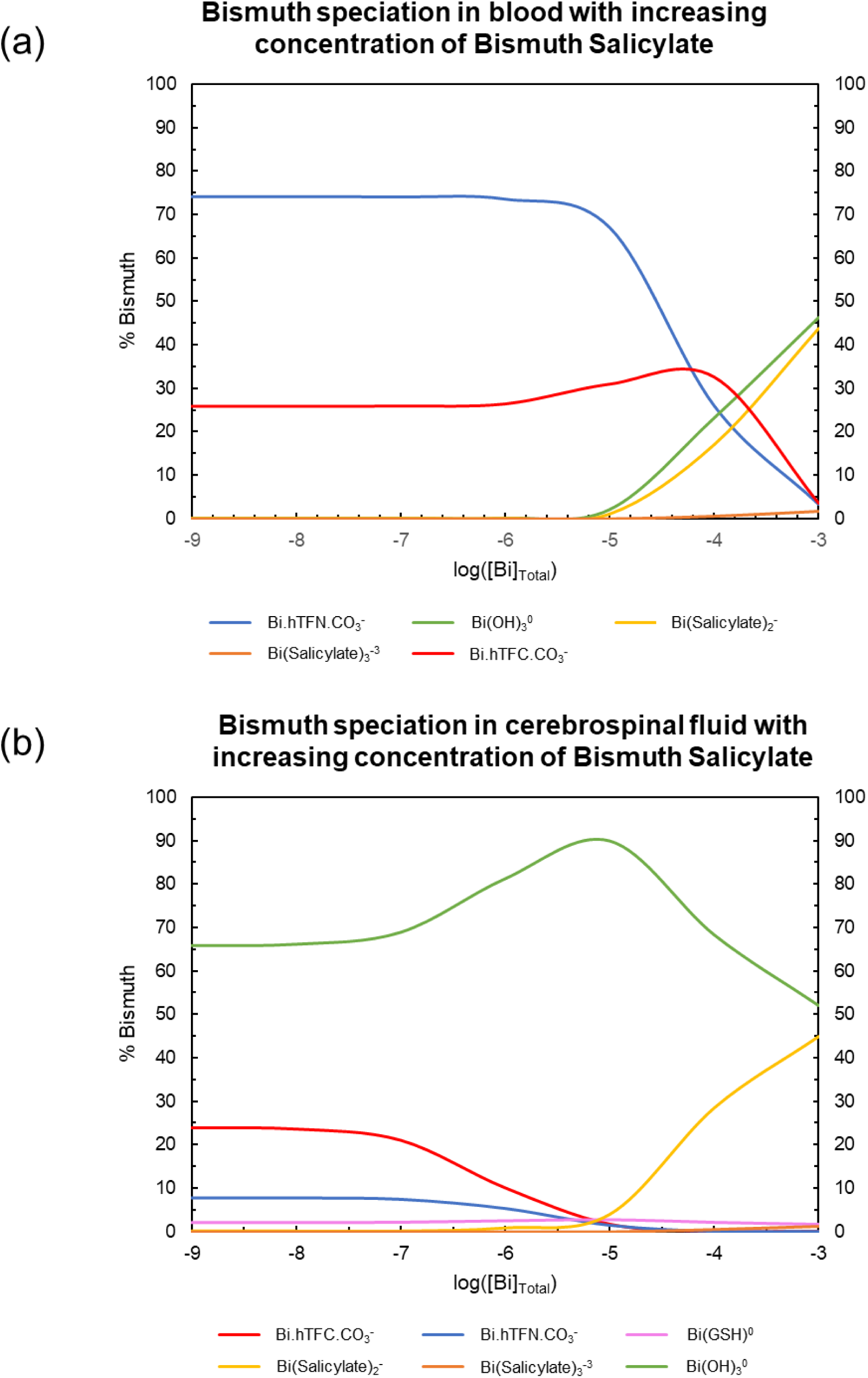
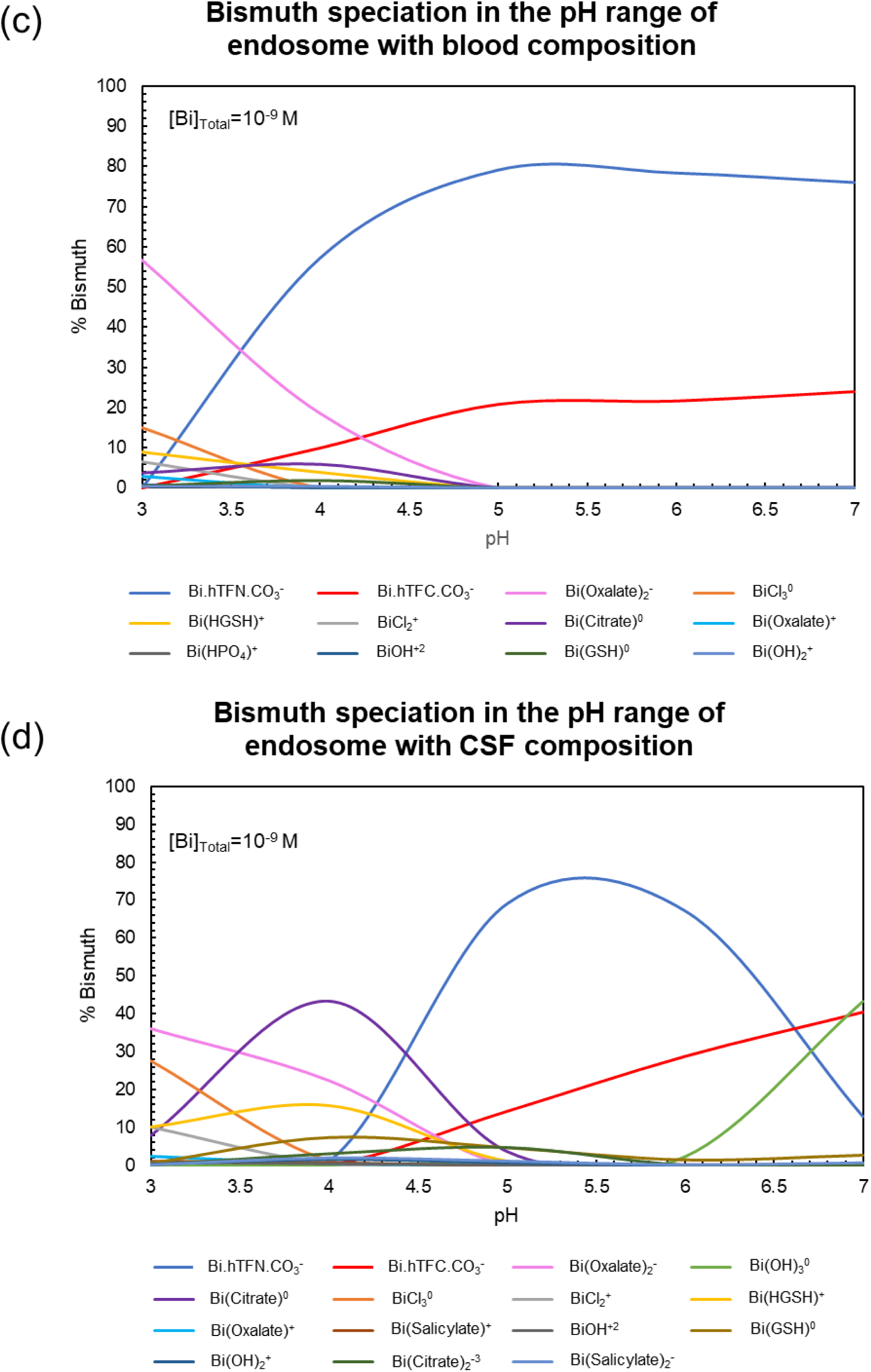
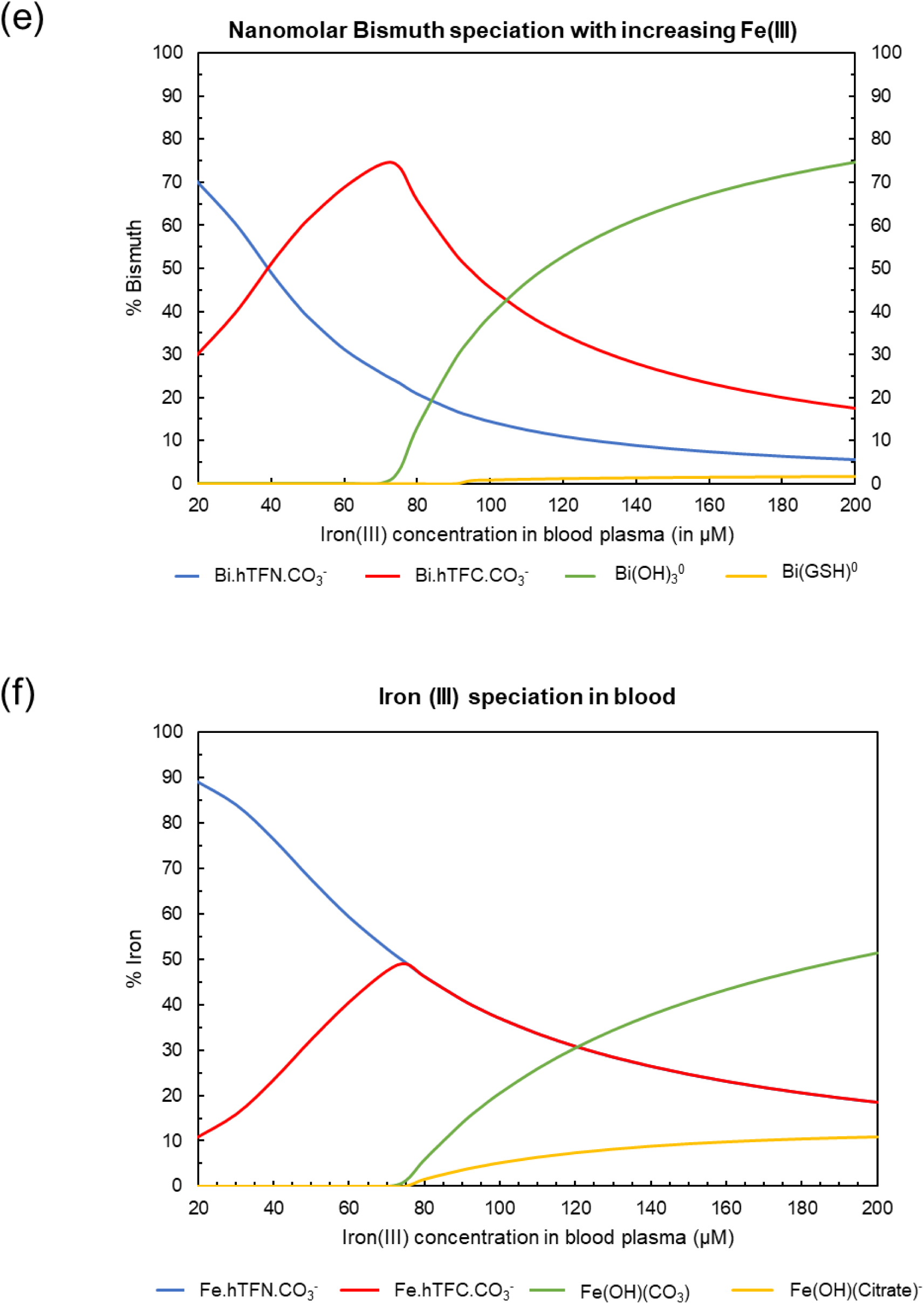
Speciation and bioavailability of bismuth in human bodily fluids.

As a considerable amount of bismuth is present in blood as the transferrin complex, it is likely to be transported to cells via receptor-mediated endocytosis (Guo et al. 2000, Zhang et al. 2001). As the endosomal pH ranges from 3.5 to 6.5 (Diering & Numata 2014, Hu et al. 2015), bismuth speciation is likely to be considerably different under these conditions than its speciation in blood plasma. Since the chemical composition of the endosome is currently unknown, we performed speciation calculations with blood and CSF compositions over the known pH range of the endosome. The results are shown in Figs. 6c and 6d, which both illustrate differences in bismuth speciation as functions of pH. At low pH, oxalate complexes constitute a considerable fraction of bismuth in either the blood or the CFS models of the endosome. In the case of CSF, Bi-citrate complexes are also predicted to be relatively abundant at low pH as shown in Fig 6d. In both cases, transferrin complexes dominate the speciation at pH >5, and it is rivaled in abundance by Bi(OH)_3_ at higher pH in the CSF model. These results are consistent with the reported release of metal (Fe^+3^ and Bi^+3^) from transferrin at pH 4.5-2.0 (Guo et al. 2000, Zhang et al. 2001) and have direct implications for the bioavailability of bismuth.

In addition to transferrin concentration and pH, we found that bismuth speciation also depends strongly on iron(III) abundance. As iron(III)-transferrin stability constants are larger than the corresponding bismuth stability constants, iron spontaneously displaces transferrin-bound bismuth, thereby increasing the bismuth-LMM fraction. The calculated dependence of total plasma iron (III) concentration on bismuth speciation is shown in Fig. 6e. Note that the transferrin fraction of bismuth maximizes at 74 µM iron (III), which corresponds to the maximum metal-carrying capacity of transferrin. Exceeding this maximum is likely to transpire in cases of genetic iron-overload (hemochromatosis). At a total iron (III) concentration of 80 µM, the non-transferrin bound iron (NTBI) concentration amounts to ∼6 µM (Fig. 6f), which is commonly seen in patients suffering from some form of iron-overload (Brissot et al. 2012). At this concentration, our calculations reveal that about 14% of bismuth exists as the neutral LMM complex Bi(OH)_3_, which may be bioavailable. Thus, iron-overload conditions may be associated with high bismuth cytotoxicity.

In summary, we find that bismuth speciation depends strongly on transferrin, proton and iron (III) abundance in the biological system of interest. Bismuth speciation is considerably different in blood compared to cerebrospinal fluid, which may help explain the associated neuropathy of the metal. Our calculations of bismuth speciation as a function of pH are consistent with experimental measurements performed in simpler systems. It may be worth reiterating that estimates of stability constants for bismuth complexes with oxalate and salicylate were crucial in obtaining these simulations. We hope these models can drive further investigation in the exciting field of bismuth-based drugs.

### 4.5 Rhodium-based anticancer drugs

Rhodium-based drugs have been of interest to the anticancer community for over six decades (Taylor & Carmichael 1953, Giraldi et al. 1977, Rao et al. 1980, Kopf-Maier 1994, Katsaros & Anagnostopoulou 2002, Junicke et al. 2003, Jungwirth et al. 2011, Domotor & Enyedy 2019). The predominant mode of action of rhodium and other platinum-group metals (like ruthenium, palladium, osmium and iridium) differs from that of gallium-based anticancer drugs. The platinum-based drugs appear to bend DNA by cross-linking adjacent guanines leading to adherence by DNA-binding proteins (Jamieson & Lippard 1999). Rhodium complexes have been found to interfere with oncogenic signaling and cancer-promoting or epigenetic regulatory processes (Leung et al. 2012, Kang et al. 2017). Most rhodium compounds eliciting anticancer activity are octahedral complexes of polypyridine, polyquinoline and other aromatic chelates. Here we report rhodium speciation in blood plasma with two chelators-maltol and oxine that are known to increase metal bioavailability as discussed above. Additionally, we report rhodium speciation with the first rhodium complex known to have anti-tumor activity-RhCl_3_ (Taylor & Carmichael 1953).

Perhaps the extremely high cost of rhodium over the last century (Zientek et al. 2017) has contributed to the limited thermodynamic data for rhodium complexes. In lieu of this, stability constants for rhodium complexes can be estimated to obtain rhodium speciation in any biological fluid system of interest. In this study, values were estimated using multiple linear free energy relationships for rhodium complexes of maltol and oxine, as well as the ligands listed in Table 1 including transferrin and albumin (MBS site) (Prasad & Shock 2025b). Additional stability constants for rhodium-chloride complexes were taken from Cozzi & Pantani (1958) while the solubility product of Rh(OH)_3_(s) was taken from Forrester & Ayres (1959).

Predicted changes in the speciation of rhodium in blood with increasing concentration of RhCl_3_ is depicted in Fig. 7a. It can be seen that almost all of the rhodium is bound to the proteins transferrin and albumin. Transferrin dominates the speciation at low rhodium abundance as it has a higher stability constant but gets saturated at about 60 µM rhodium as its carrying capacity is reached. At this point, the albumin fraction of rhodium begins to increase with increasing total RhCl_3_ concentration and dominates the speciation at ≥ 120 µM rhodium. Note that at ∼70 µM rhodium, Rh(OH)_3_(s) reaches saturation.

**Fig 7(a-c).**
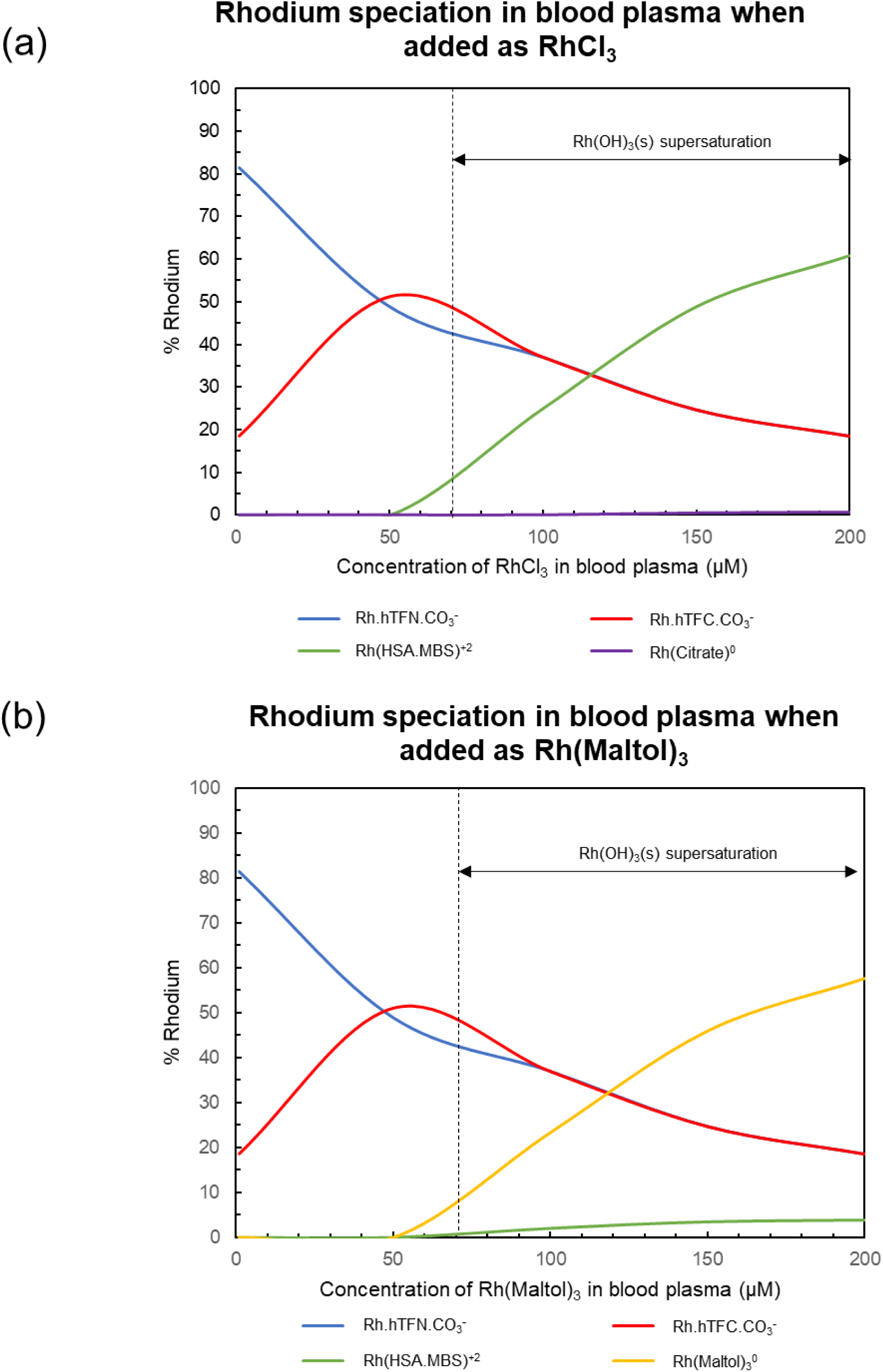
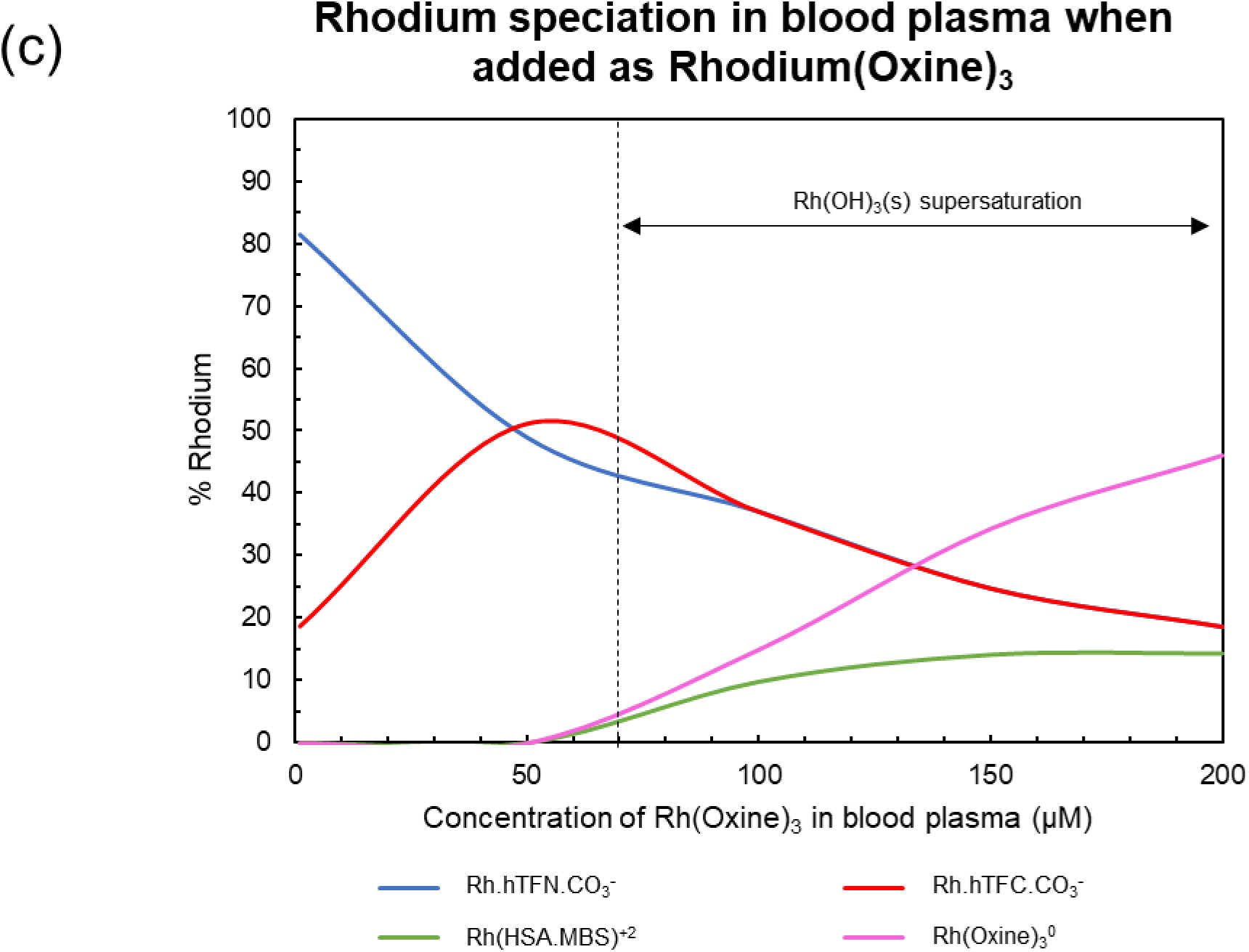
Rhodium speciation in blood plasma.

Similar calculations with Rh(Maltol)_3_ and Rh(Oxine)_3_ are displayed in Fig. 7b and 7c, respectively. While rhodium speciation is virtually identical in both cases up to ∼50 µM rhodium, it differs considerably beyond this concentration. At 200 µM of total Rh(Maltol)_3_, about 58% of the rhodium exists as Rh(Maltol)_3_^0^ while at the corresponding concentration of total Rh(Oxine)_3_, ∼45% of rhodium exists as Rh(Oxine)_3_^0^. However, at 100 µM, the distribution of rhodium as the corresponding tris complexes (1:3 metal-ligand complex) is about the same at ∼20%, which corresponds to a considerable bioavailable fraction assuming no losses from precipitation.

We hope these results outline the practical utility of performing speciation calculations for medical applications involving expensive metals like rhodium. Similar calculations can prove to be useful for other organometallic forms of rhodium that are being used for pharmaceutical applications (Öhrström 2006, Domotor et al. 2014, Domotor & Enyedy 2019). As these are mixed-ligand complexes, the strategy for stability constant estimation is likely to differ from methods used in this study. We hope such theoretical endeavors are mutually assisted by progress in stability constant measurements and elucidation of rhodium-associated cytotoxic activity. As rhodium has several other applications including in catalytic converters of automobile exhaust and in the manufacture of nitric oxide that is the raw material for explosives, fertilizers, and nitric acid (Öhrström 2006 and Zientek et al. 2017), it is a relevant metal todat and is likely to stay pertinent in the near future (Jasinski et al. 2018).

## 5. Conclusion

Previously, a shortcoming of metal speciation modeling in biological systems was the limited number of relevant metal complexes, especially involving organic compounds (Kiss et al. 2017, Wilke et al. 2017). We ameliorated this problem by estimating stability constants of over 18,000 metal-organic complexes including proteins, peptides, amino acids, carboxylates and phenols (Prasad & Shock 2025a and Prasad & Shock 2025b). In this work, we used those estimates to conduct metal speciation calculations of 10 metals in blood plasma, a biological system integral to human health. As a foundation, we predicted the speciation of the seven predominant metals in blood. In addition, we calculated the speciation of gallium and rhodium based anticancer drugs, copper in Wilson Disease, zinc-based antidiabetics and bismuth-based antacids. A special emphasis was placed on the implications of our speciation calculations for the bioavailability of these metals and the associated therapeutic or toxic potential.

Our calculations reveal that proteins dominate the speciation of all metals and are therefore crucial for model construction of metal transport and distribution in the human body. Additionally, proteins have distinct functional groups that select for specific metals and mirror the selectivity seen for LMM ligands. Other predominant factors that significantly affect metal speciation are pH, chelator concentration and competition with other metal ions. In addition to blood plasma, we predicted metal speciation for cerebrospinal fluid and for associated vesicles at low pH.

A major part of our focus was to limit errors in speciation models, which allows us to identify areas where improvements can be made. Based on our assessment, care is warranted while determining stability constants for metal-protein complexes. Currently, these values are obtained via competition experiments with LMM ligands like EDTA and NTA. Consequently, metal-protein log K values are extremely sensitive to the corresponding property associated with the metal complexes of these competitive ligands. Thus, accurate stability constants of these complexes, including all multi-ligand and protonated/hydroxylated forms, are needed to obtain accurate values of metal-protein log K. Additionally, several studies on metal-protein equilibria do not report and may not monitor the ionic strength of the system. Improvements for metal-LMM complexes can also be made, especially for mixed ligands of biological relevance like compounds associated with platinum-based anticancer compounds.

Besides the physiological conditions discussed in this paper, metals are involved in aging (Bredesen 2015), Alzheimer’s Disease (Adlard & Bush 2018), manganese toxicity (Michalke et al. 2017), lead toxicity (Yedju et al. 2010) and lithium-based antidepressants (Cipriani et al. 2013). Additionally, metals like gadolinium, cobalt and technetium are used in diagnostic medical imaging as radioisotopes or contrast agents (Jackson & Byrne 1996 and McInnes et al. 2017). In this work, we have tried to bridge the gap between the biotic and abiotic aspects of metal speciation using the foundational theory of thermodynamics. We hope this work helps induce a more symbiotic association between the fields of metal-speciation modeling and metal-based pharmacology. We firmly believe that both fields not only have a lot to offer each other but each can drive innovation in its counterpart.

## Acknowledgements

This work was supported by Deep Carbon Observatory. The authors would like to thank Arizona State University (ASU) and all the past and current members of Group Exploring Organic Processes In Geochemistry (GEOPIG) at ASU for helpful discussions.

## Appendix: Procurement of metal-protein stability constants via regression of metal speciation measurements from metal-protein binding studies

We validated our metal-protein stability constant data by regressing metal-speciation measurements of studies reporting metal-protein binding constants. This was primarily performed for the proteins human serum transferrin (hTF) and human serum albumin (HSA) as these are the two predominant proteins found in blood plasma. This regression was crucial for transferrin as several studies report pH-dependent ‘stoichiometric constants’ that deviates from pH-independent equilibrium constants used in our models. Additionally, many of these metal-binding studies for transferrin were performed before the measurement of the Tyr 188 pK_a_ at the binding site that was subsequently found to be considerably different from the pK_a_ of free tyrosine side chain (Sun et al. 2004). Incorporation of this pK_a_ value in our calculations enabled regression of experimental data with a pH-independent stability constant for several metals.

Equilibrium dialysis experiments of Aisen et al. 1978 were simulated to obtain stability constants of Fe(III)-Transferrin. In these experiments, a semi permeable membrane was used to constrain the large protein molecule to the ‘protein compartment’ followed by measurement of total iron concentration in the ‘protein compartment’ and the ‘buffer compartment’. As the free ferric ion (Fe^+3^) could move freely across this membrane, its concentration would be expected to be the same in ‘protein’ and ‘buffer’ compartments. As may be seen from Fig. A1, this was found to be the case for most experiments carried out at two different pH. The deviations from the 1:1 line occur at ferric ion molality which can be attributed to the higher measurement uncertainty associated with low concentration. A set of pH-independent stability constant values were obtained (logK_1_= 18.0, logK_2_= 16.7) that were different from the pH dependent values reported in the original study (logK_1_*= 19.5 & logK_2_*= 17.4 at pH=6.7 and logK_1_*= 20.7 & logK_2_*= 19.4 at pH=7.4).

**Fig. A1.**
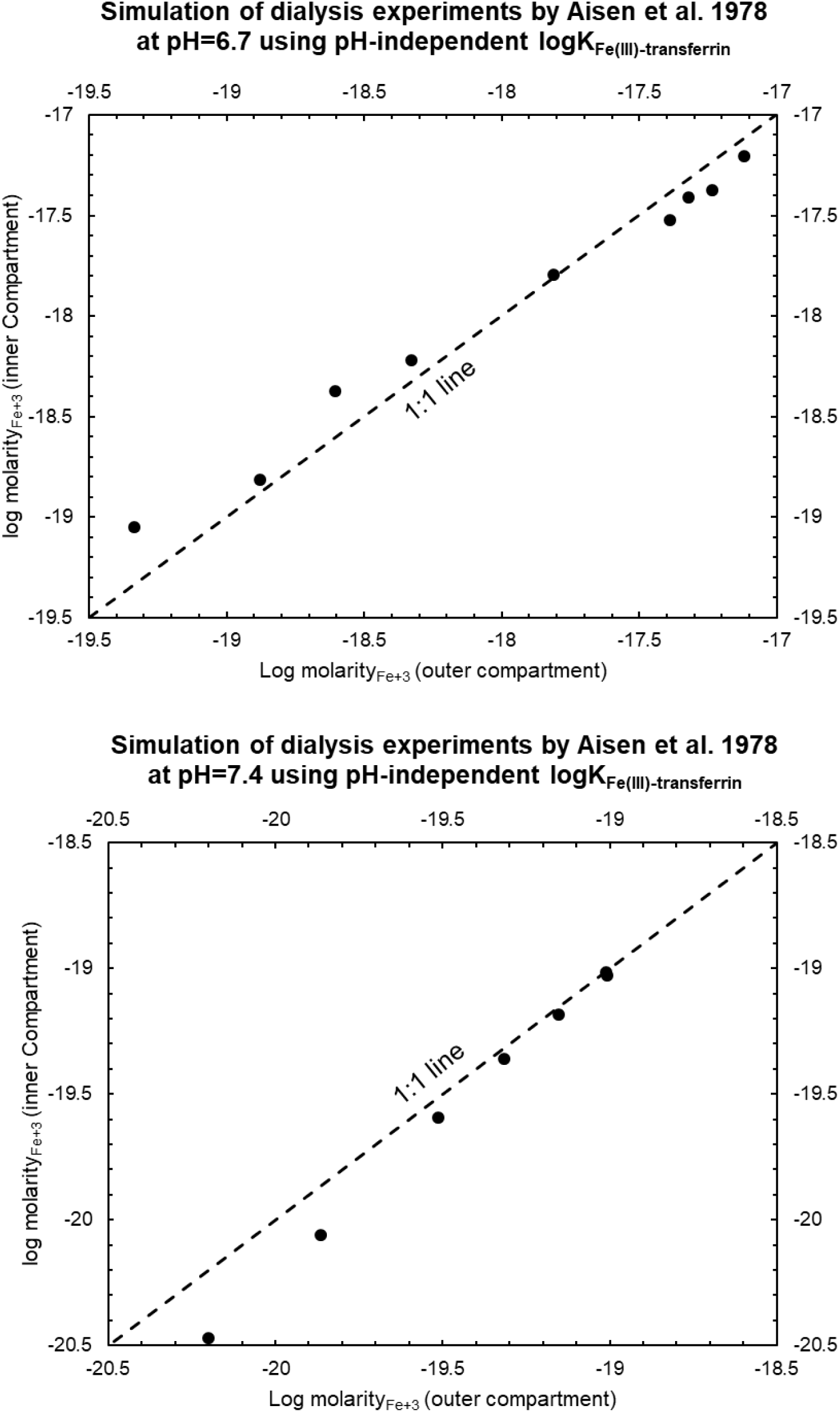
Analysis of dialysis experiments by Aisen et al. 1978 using pH-independent logK_Fe(III)-transferrin_ and subsequently published results on Tyr 188 pK_a_ by Sun et al. 2004.

Analogous regression was performed for transferrin binding studies with Zn^+2^, Bi^+3^ and Ga^+3^ and are shown in Fig. A2. In these studies, a metal chelator was used as a competitive ligand to control the abundance of metal-transferrin complex that was measured using UV or NMR spectroscopy. The standard state stability constant obtained were slightly different from the stoichiometric constants dependent on pH and concentration of bicarbonate (logK_1_= 4.0, logK_2_= 3.0 vs. logK_1_*= 7.9 & logK_2_*= 6.5 for Zn^+2^; logK_1_= 17.2, logK_2_= 16.4 vs. logK_1_*= 20.1 & logK_2_*= 19.3 for Bi^+3^; logK_1_= 17.5, logK_2_= 16.2 vs. logK_1_*= 20.4 & logK *= 19.4 for Ga^+3^).

Unexpectedly, ligand competition studies for metal-albumin studies were extremely rare (Kiss et al. 2017). Out of the 10 metal ions studied in this work, we only found ligand competition studies for Zn(II)-albumin binding wherein dipicolinate was used as the competing ligand (Bytzek et al. 2009). Zinc speciation was measured using capillary zone electrophoresis-inductively coupled plasma-mass spectrometry (CZE-ICP-MS). In this case, Zn-Albumin stability constants independently reported by Lu et al. 2012 and other relevant stability constants measured by the group (Kiss et al. 2009 and Enyedy et al. 2008) replicated the original experiments within 10% of zinc at high Zn-HSA concentration and within 15% zinc at low Zn-HSA abundance (Fig. A3).

**Fig. A2.**
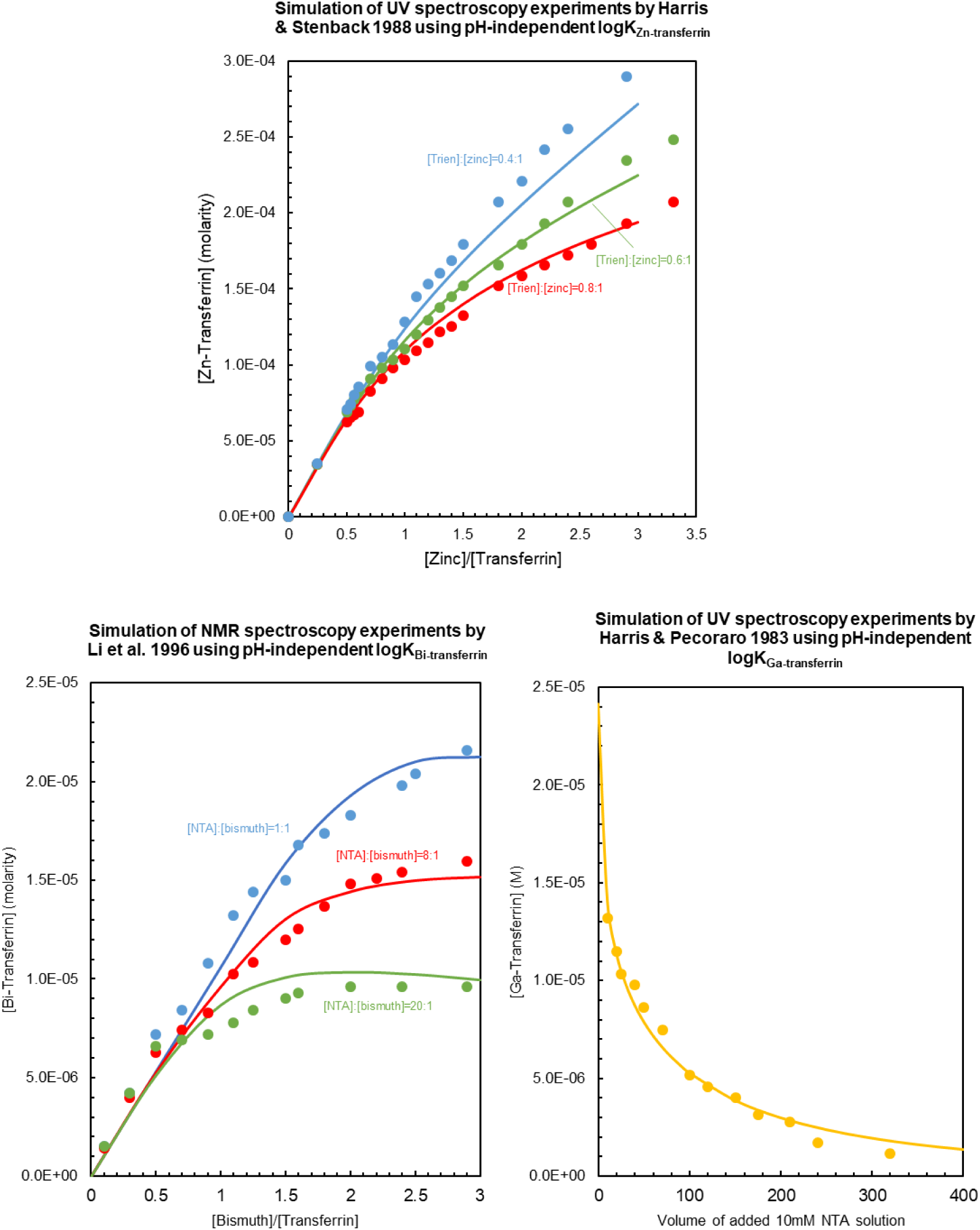
Analysis of UV and NMR spectroscopy measurements used to determine transferrin stability constants with zinc, bismuth and gallium performed by Harris &. Stenback 1988, Li et al. 1996 and Harris & Pecoraro 1983.

**Fig. A3.**
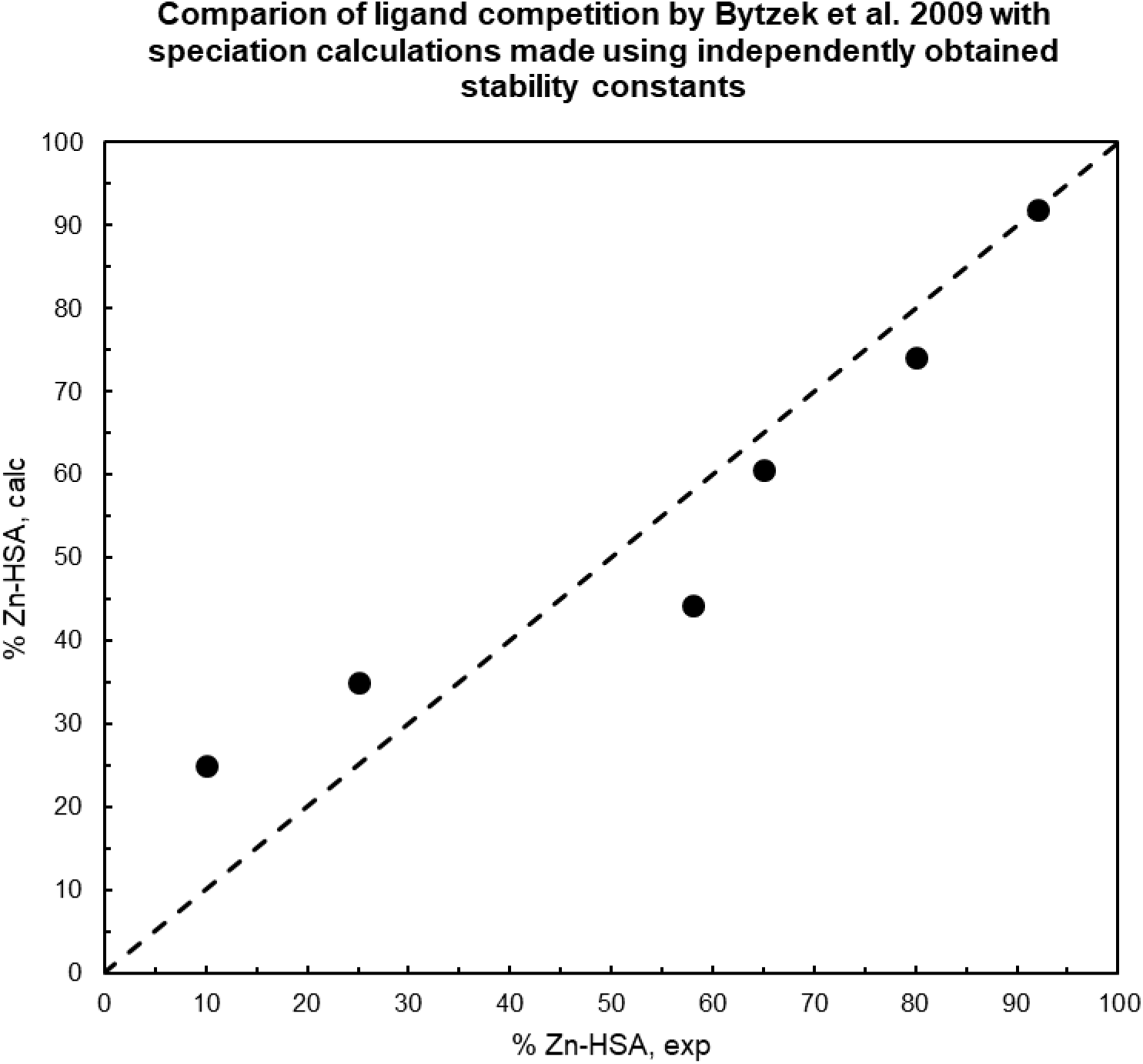
Analysis of CZE-ICP-MS (capillary zone electrophoresis-inductively coupled plasma-mass spectrometry) experiments by Bytzek et al. 2009 in zinc-human serum albumin-dipicolinate system with independently obtained stability constants. The logK_Zn-HSA_ used here was used in speciation calculations in this work.

